# Guiding the design of bacterial signaling interactions using a coevolutionary landscape

**DOI:** 10.1101/116947

**Authors:** R. R. Cheng, E. Haglund, N. Tiee, F. Morcos, H. Levine, J. A. Adams, P. A. Jennings, J. N. Onuchic

## Abstract

The selection of amino acid identities that encode new interactions between two-component signaling (TCS) proteins remains a significant challenge. Recent work constructed a co-evolutionary landscape that can be used to select mutations to maintain signal transfer interactions between *partner* TCS proteins without introducing signal transfer between non-partners (crosstalk). A bigger challenge is to introduce mutations between non-natural partner TCS proteins using the landscape to enhance, suppress, or have a neutral effect on their basal signal transfer rates. This study focuses on the selection of mutations to a response regulator (RR) from *Bacilus subtilis* and its effect on phosphotransfer with a histidine kinase (HK) from *Escherichia Coli*. Twelve single-point mutations of the RR protein are selected from the landscape and experimentally expressed to directly test the theoretical predictions on the effect of signal transfer. Differential Scanning Calorimetry is used to monitor any protein stability effects caused by the mutations, which could be detrimental to proper protein function. Of these proteins, seven mutants successfully perturb phosphoryl transfer activity in the computationally predicted manner between the TCS proteins. Furthermore, brute-force exhaustive mutagenesis approaches indicate that only 1% of mutations result in enhanced activity. In comparison, of the six mutations predicted to enhance phosphotransfer, two mutations exhibit a significant enhancement while two mutations are comparable to the wild-type. Thus co-evolutionary
landscape theory offers significant improvement over traditional large-scale mutational studies in the efficiency of selecting mutations for protein engineering and design.

## Introduction

Bacterial two-component signaling (TCS) (Capra & Laub, 2012, Casino, Rubio et al., 2010, Hoch, 2000, Laub & Goulian, 2007, Stock, Robinson et al., 2000, Szurmant & Hoch, 2010) is the primary means by which bacteria respond to external stimuli. TCS is carried out by two partner proteins working in tandem: a histidine kinase (HK) and a response regulator (RR). The HK detects a stimulus and proceeds to autophosphorylate, after which its partner RR can bind to it and receive the phosphoryl group. The phosphorylated RR can then function as a transcription factor that regulates gene expression. Although there are as many as 10^2^-10^3^ homologous TCS pairs in a bacterium, each controlling the response to a different stimulus, a HK typically transfers signal with its partner RR. This preference between partners is facilitated by residue interactions between the HK and RR, which form their mutual binding interface.

Selecting mutations that encode new interactions between non-partner TCS proteins remains a significant challenge in synthetic biology despite recent successes via extensive mutagenesis and selection. Mutagenesis studies have demonstrated that TCS interactions can be engineered through amino acid mutations *in vitro* (*Capra, Perchuk et al., 2010, Skerker, Perchuk et al., 2008*) and *in vivo* (*Skerker et al., 2008*). Similar mutagenesis approaches have been used to restore autophosphorylation in a chimeric HK (Dago, Schug et al., 2012), which is composed of a dimerization domain and an ATPase domain from different HK proteins. Recently, a comprehensive mutational study (Podgornaia & Laub, 2015) explored the full sequence-space of 4 residues on a HK that still maintained functional signal transduction with its partner RR *in vivo*. While it is clear that the technology to exhaustively mutate and interrogate the functionality of hundreds of thousands of mutations *in vitro/vivo* exists, it is not feasible to design new TCS interactions by experimentally constructing all amino acid possibilities. Therefore, the design of new TCS interactions would greatly benefit from computational, data-driven approaches, which learn from the amino acid combinations that have been naturally selected.

Recent data-driven approaches(Ekeberg, Lovkvist et al., 2013, Morcos, Pagnani et al., 2011) have advanced the quantitative modeling of protein sequence data. These approaches have been used to quantify the correlated mutational patterns that arise from the constraint to maintain residue-residue interactions during natural selection(de Juan, Pazos et al., 2013), referred to as amino acid coevolution. Highly coevolving residue pairs have been shown to be contacts in a 3D protein structure or complex(Ekeberg et al., 2013, Marks, Hopf et al., 2012, Morcos et al., 2011, Ovchinnikov, Park et al., 2017). These approaches have also been used to investigate mutational landscapes, which quantify how amino acid mutations affect organism phenotypes(Cheng, Nordesjo et al., 2016, Figliuzzi, Jacquier et al., 2016, Hopf, Ingraham et al., 2017). In particular, a previous study focused on the interprotein coevolution between the HK and RR protein residues to construct a mutational landscape, 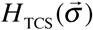 (Eq. 1), which can be calculated as a proxy for signal transfer efficiency for an arbitrary pair of TCS proteins with sequence, 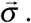 Mutational effects on signaling can then be calculated as 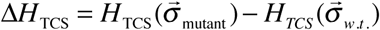 between a mutant sequence, 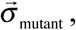 and a wild-type sequence, 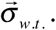 Mutations that enhance, suppress, or are neutral to signaling are reflected by the conditions Δ*H*_TCS_ < 0, Δ*H*_TCS_ > 0, and Δ*H*_TCS_ ≈ 0, respectively.

As Δ*H*_TCS_ can be used to identify mutations in a TCS protein that maintained functional signaling with its native partner (Cheng et al., 2016), consistent with the findings from *in vivo* experiment (Podgornaia & Laub, 2015), the expansion of this approach toward non-native TCS proteins is necessary to test its predictions. The current study expands upon this idea of selecting mutations directly from 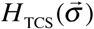 to rationally encode new TCS interactions between a HK and a RR protein that are not native partners, henceforth called non-partners. Specifically, mutations are selected for the RR protein, Spo0F from *Bacillus subtilis*, to enhance its *in vitro* signal transfer with a non-partner HK protein, EnvZ from *Escherichia coli*. Mutations are also selected from the landscape to suppress or have a neutral effect on the phosphotransfer between EnvZ and Spo0F. To test the predictions of the inferred model, the phosphotransfer between the mutant Spo0F and EnvZ is measured *in vitro* using a protein radiolabeling assay similar to that used for KinA/Spo0F phosphotransfer *in vitro* (Tzeng & Hoch, 1997). This analysis is combined with *in vitro* experiments using Differential Scanning Calorimetry (DSC) to measure the effect of each specific mutation on the enthalpy of protein unfolding, ΔΔ*H*_DSC_, to ascertain the mutational effect on folding and stability. The effect of mutations to Spo0F on other HK homologs (*in vivo* cross-talk) is not considered in this study, but can also be successfully recapitulated by the model, which is discussed in great detail in a recent theoretical study (Cheng et al., 2016).

The results of this study are organized in three subsections: (*i*) The selection of mutations from the coevolutionary landscape, (*ii*) The comparison of the landscape predictions with the experimentally measured phosphotransfer between Spo0F and EnvZ, and (*iii*) The comparison of the computational predictions on Spo0F stability and the experimental finds using DSC.

## Results

### Inferring candidate mutations for the RR Spo0F

Candidate mutations for Spo0F to encode its preferential interaction with the HK EnvZ are selected from the subset of mutations for which 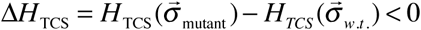 (Eq. 1), where 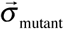 and 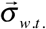 are the mutant and wild-type sequences, respectively. Such mutations can be interpreted as increasing the signal transfer efficiency (Cheng et al., 2016) between EnvZ and Spo0F, according to the inferred quantitative model.

In this study, the selection of mutations is limited to single residue sites on Spo0F that (*i*) form contacts with the HK in the representative HK/RR complex and that are also (*ii*) coevolving with the residues of the HK. Assuming that the HK/RR binding interface is preserved over the majority of TCS partners, the predicted KinA/Spo0F complex (Cheng, Morcos et al., 2014) is used as a representative structure (Figure 1). In *B. subtilis*, KinA is the HK that phosphorylates Spo0F in the sporulation phosphorelay (Burbulys, Trach et al., 1991). The predicted complex, used in this study, is consistent with an earlier predicted TCS complex (Schug, Weigt et al., 2009) as well as an experimental crystal structure (Casino, Rubio et al., 2009). Figure 1A shows the number of contacts, *N*_*contact*_, formed between Spo0F and KinA in the representative structure using a 10Å cutoff between all heavy atoms. Four main groups of residues on Spo0F form the contacts with the HK and are mapped to the sequence and secondary structure (1C), i.e., α1 (Group 1), β3-α3 loop (Group 2), β4/β4-α4 loop (Group 3), and β5-α5 loop/α5 (Group 4).

**Figure 1.**
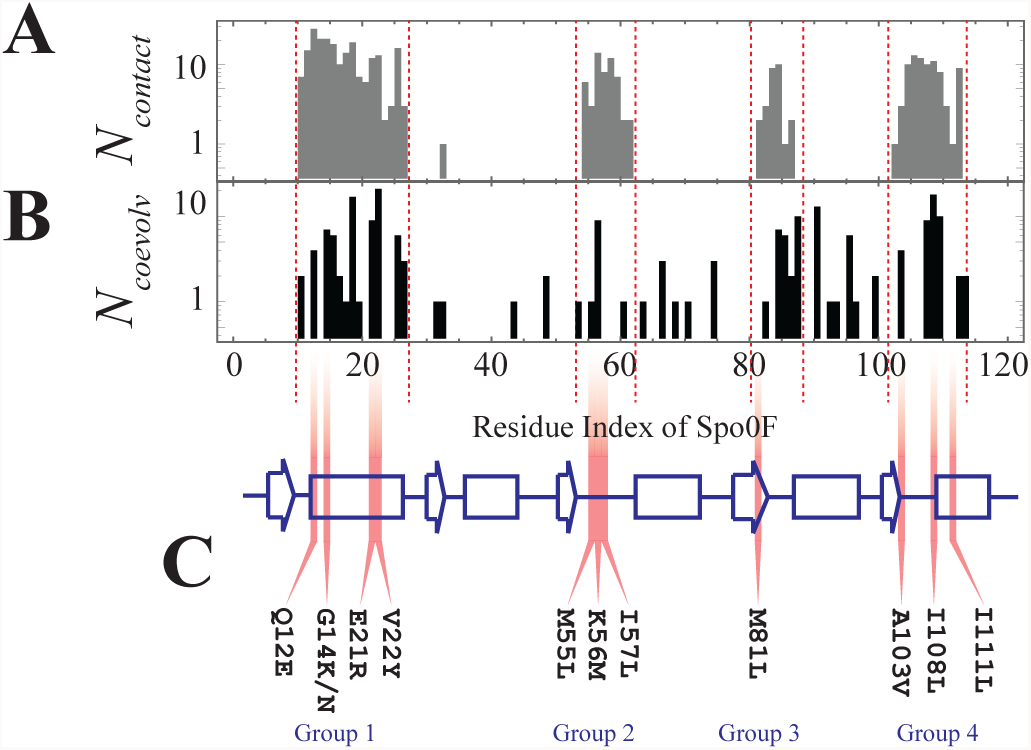
Selecting mutants from highly coevolved regions of the HK/RR interface. (A) A histogram of the number of contacts, *N*_*contact*_, formed between the RR Spo0F and the HK KinA in a representative structure of the TCS complex is plotted as a function residue number on Spo0F (Cheng et al., 2014). (B) Using coevolutionary analysis of HK/RR partner sequences, the top 200 coevolving HK/RR interprotein residue pairs are calculated using Direct Information (DI) (Eq. 3). For these top coevolving residue pairs, the number of HK residues coevolving with each RR residue, *N*_*coevolv*_, is plotted in a histogram as a function of the residue numbers of the RR protein family, which are mapped on to the corresponding residue numbers on Spo0F. (C) The secondary structure of Spo0F is drawn as a cartoon, with strands denoted by arrows, helices by rectangles and loops and turns denoted by lines. Mutations in highly coevolving RR residues that formed contacts with the HK in this study were obtained from the four groups: α1 (Group 1), β3-α3 loop (Group 2), α4/α4-α4 loop (Group 3), and β5-α5 loop/α5 (Group 4).

The amount of coevolution between all interprotein residue pairs (i.e., HK/RR residue pairs) according to the inferred model is quantified using the Direct Information (DI) (Eq. 3) (Morcos et al., 2011, Weigt, White et al., 2009). The coevolving interprotein residue pairs (between the HK and RR) form contacts that stabilize the TCS complex (Cheng et al., 2014, Schug et al., 2009, Weigt et al., 2009). Figure 1B shows the number of HK residues found to strongly coevolve with each RR residue, *N*_*coevolv*_. Spo0F residue positions with a high *N*_*coevolv*_ are interpreted as being candidate sites for encoding new TCS interactions.

The primary candidates for enhancing the phosphotransfer between EnvZ and Spo0F are thus chosen from the overlap between Figures 1A and 1B, for mutations that satisfy Δ*H*_TCS_ < 0 (Eq. 1). These primary candidates are G14K, E21R, and V22Y from Group 1, M55L from Group 2, and I108L and I111L from Group 4. Additional mutations are also selected from the four contact groups. The mutations G14N and K56M are chosen because they are predicted to be highly deleterious to phosphotransfer between EnvZ and Spo0F (Figure 2A), i.e., Δ*H*_TCS_ > 0. The remaining mutations that are also explored include Q12E, I57L, M81L, and A103V, which are predicted to have a neutral effect on the phosphotransfer between EnvZ and Spo0F. The computational predictions of the signal transfer efficiency, Δ*H*_TCS_, for all mutants are shown in Figure 2A. Figure 2B shows the experimental results, which are discussed in the proceeding subsection (vida supra). Figure 2C shows all of the single-site mutations plotted together on the representative structure of Spo0F. All of these mutational sites physically interact with the HK in the representative structure of a TCS complex (Figure 2D).

**Figure 2.**
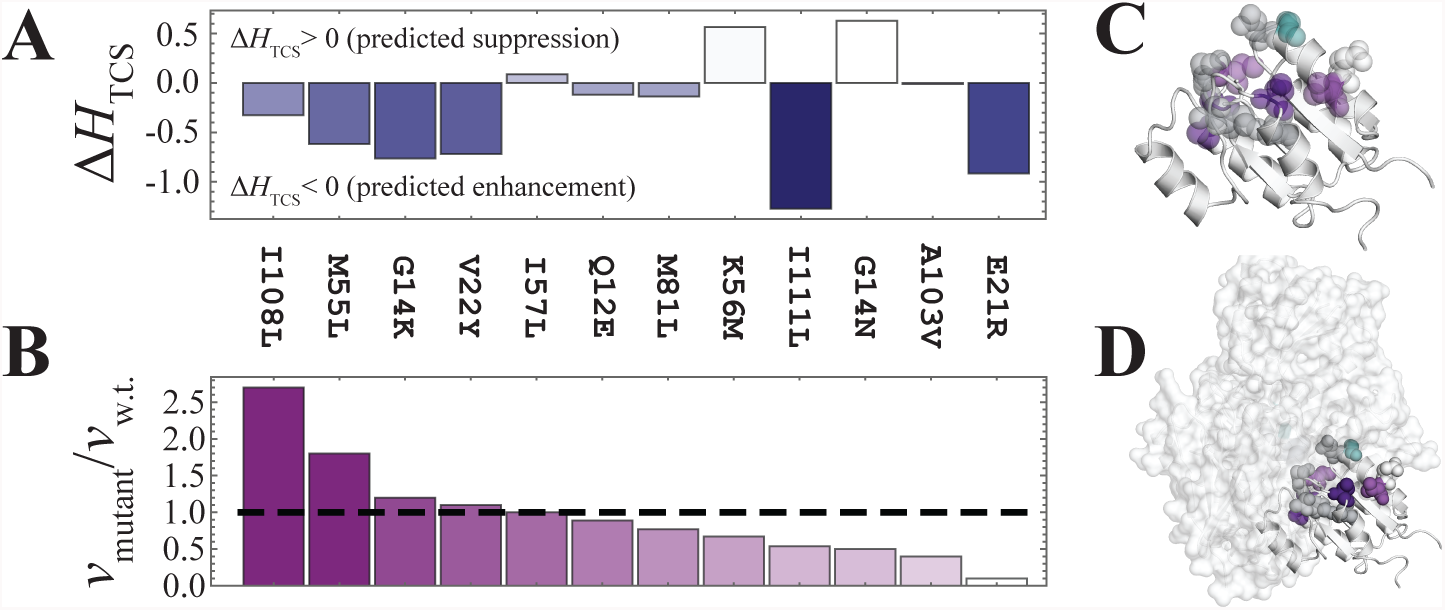
Comparison of mutational predictions of Δ*H*_TCS_ with *in vitro* phosphotransfer rates. (A) The computational proxy for signal transfer efficiency, Δ*H*_TCS_ (Eq. 1), is plotted for each of the Spo0F mutations. By definition, Δ*H*_TCS_ < 0 represent mutations that are predicted to enhance phosphotransfer between EnvZ and Spo0F, while Δ*H*_TCS_ > 0 represent mutations that suppress phosphotransfer. (B) The *in vitro* phosphotransfer rate between EnvZ and each of the Spo0F mutants, *v*_mutant_, is plotted normalized by the phosphotransfer rate between wild type EnvZ/Spo0F, *v*_w.t._. Here, *v*_mutant_ / *v*_w.t._ > 1 shows mutations that enhanced the phosphotransfer rate. Four out of these six mutants were able to enhance phosphotransfer, showing the predictive capabilities of the coevolutionary model. Neutral and deleterious mutations were also successful, as discussed below. (C) The mutations (colored by group) are plotted on the representative structure of Spo0F (PDB ID: 1PEY) (Mukhopadhyay, Sen et al., 2004). (D) The mutations (colored in purple and represented as spheres) are shown on Spo0F bound to KinA in the representative TCS complex (Cheng et al., 2014). The mutated residues are colored to match (B), where a darker purple represents a greater enhancement to the experimental phosphotransfer rate. The G14K/N mutations are shown in cyan. The structural representations in (C) and (D) were generated using PyMOL (Schrodinger, 2015).

### Comparison of phosphotransfer predictions with *in vitro* experiment

Figure 2A shows the predicted effect of mutations to Spo0F on the EnvZ/Spo0F phosphotransfer, where mutants with Δ*H*_TCS_ < 0 are predicted to exhibit a phosphotransfer enhancement with respect to the wild-type EnvZ/Spo0F interaction. The mutants G14K, E21R, V22Y, M55L, I108L and I111L are predicted to enhance phosphotransfer, whereas G14N and K56M are expected to decrease phosphotransfer. The remaining mutations are predicted to have a neutral affect on the EnvZ/Spo0F interaction.

The phosphotransfer reaction between EnvZ and Spo0F (wild-type and mutants) was measured *in vitro* to obtain phosphotransfer rates for wild-type and mutant Spo0F, denoted as *v*_w.t._ and *v*_mutant_, respectively. The ratio of the mutant phosphotransfer rate to the wild-type phosphotransfer rate, *v*_mutant_ / *v*_w.t._, is shown in Figure 2B. Of the 6 mutants predicted to enhance EnvZ/Spo0F phosphotransfer, M55L and I108L exhibited significant enhancement, G14K and V22Y maintained wild-type activity, and E21R and I111L exhibited a significant decrease in phosphotransfer. A previous exhaustive mutagenesis study (Podgornaia & Laub, 2015) found that most amino acid mutations (∼99%) are deleterious toward TCS signaling *in vivo*. While 2 out of 6 mutations significantly enhanced phosphotransfer, 4 out of 6 mutations retained or improved wild-type phosphotransfer activity. For the two mutants predicted to suppress phosphotransfer, G14N and K56M, both decreased the wild-type EnvZ/Spo0F phosphotransfer rate, i.e., *v*_mutant_ / *v*_w.t._ < 1. Of the mutations predicted to be neutral, I57L, Q12E, and M81L exhibited a phosphotransfer rate comparable to that of the wild-type. The remaining neutral mutations were found to decrease phosphotransfer. With the exception of A103V, all mutations predicted to have a neutral effect were less deleterious to signal transfer than G14N and K56M.

### Mutational effects on protein stability measured using Differential Scanning Calorimetry (DSC)

DSC is used to measure the enthalpy of unfolding, Δ*H*_DSC_, for the wild-type and mutational variants of Spo0F. The DSC data is available as Table S1. Figure 3 shows the mutational change in the enthalpy of unfolding, i.e., 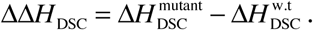 Many of the mutations found to result in a reduced phosphotransfer between EnvZ and Spo0F (Figure 2B) appear to be destabilized (i.e., ΔΔ*H*_DSC_ < 0). The loss of folding stability offers an explanation as to why several of the mutations predicted to enhance or have a neutral effect on phosphotransfer were instead found to decrease *in vitro* phosphotransfer. In particular, the mutants E21R and I111L were the two most destabilized Spo0F mutants measured by DSC (Figure 3). The destabilization of Spo0F potentially explains why both mutations, which were predicted to enhance phosphotransfer (Figure 2A), were both found to decrease phosphotransfer in experimental studies (Figure 2B).

**Figure 3.**
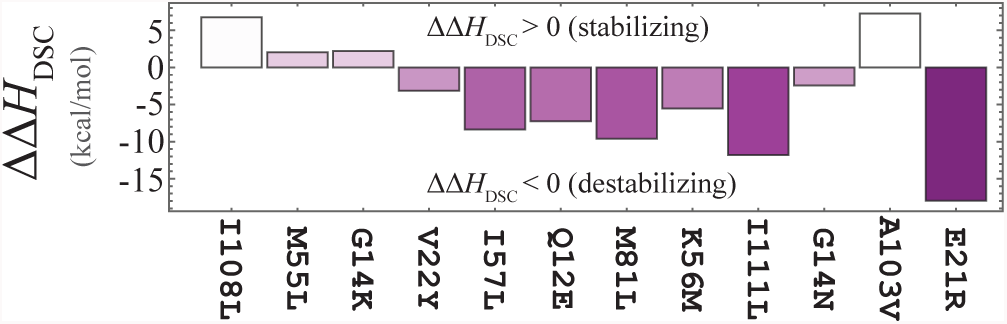
The measured mutational change in the enthalpy of unfolding. The mutational change in the enthalpy of folding, 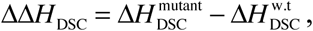 is measured using DSC for each of the Spo0F mutants with respect to the wild-type protein. Stabilizing and destabilizing mutations to Spo0F are represented by ΔΔ*H*_DSC_ > 0 and ΔΔ*H*_DSC_ < 0, respectively.

## Discussion

TCS partners have evolved to maintain interaction specificity, which is encoded in the residues that form the interface between HK and RR proteins. Selecting mutations to engineer new interactions between non-partner HK and RR proteins has remained a significant challenge in synthetic biology. The current work selects mutations directly from a co-evolutionary landscape, Δ*H*_TCS_, which serves as a proxy for signal transfer efficiency between a HK and RR (Cheng et al., 2016). Specifically, mutations are selected to enhance the signal transfer between the RR Spo0F from *B. subtilis* and the HK EnvZ from *E. coli*. These results show that 2 of the 6 mutations predicted to enhance EnvZ/Spo0F signal transfer exhibited significant enhancement *in vitro*. While it is possible to generate and interrogate the functionality of hundreds of thousands of mutations *in vivo* or *in vitro* (Podgornaia & Laub, 2015), the design of new TCS interactions can be made more practical using data-driven, computational approaches. Due to the low computational cost of generating predictions using Δ*H*_TCS_, one readily can search sequence-space for amino acid combinations that enhance signal transfer between non-partners. This combined computational and experimental approach would complement existing strategies for engineering bacterial responses that are based on modular design (Ganesh, Ravikumar et al., 2013, Hansen & Benenson, 2016, Schmid, Sheth et al., 2014, Tabor, Levskaya et al., 2011, Whitaker, Davis et al., 2012).

It was previously demonstrated by exhaustive mutagenesis that roughly 1% of mutations to the binding interface of a HK led to functional signaling in *E. coli* (Podgornaia & Laub, 2015). While not all mutations predicted to enhance *in vitro* phosphotransfer would be expected to also generate interaction specificity in living systems, 1% could serve as a rough estimate to the expected fraction of mutations that lead to phosphotransfer enhancement from a random sample. Because the mutational sequence space for any given protein is astronomically large, deciphering which mutations will improve the activity is a daunting challenge. For the 6 mutations expected to enhance activity, 2 mutations significantly improved activity. Randomly selecting 6 mutations to enhance signal transfer and generating 2 positive predictions yields a p-value of 10^-3^ using a binomial distribution, assuming that only 1% of mutations would enhance phosphotransfer. For comparison, it would take approximately 200 randomly selected mutations to obtain 2 successes on average. Moreover, as the 1% estimate applies to a random search for functional mutations (99% will be deleterious), the predictive strength of the methodology increases substantially; 4 of the 6 mutations retain or improve the wild-type EnvZ/Spo0F phosphotransfer activity. Though identifying mutations that greatly improve activity is considerably more challenging, simply assuming their statistical rarity, the methodology was also adept at determining neutral and deleterious mutations (5 of the 6 mutations correlate well with predictions). Thus, the selection of mutations directly from a TCS co-evolutionary landscape offers a truly significant enhancement in the positive predictive value compared to design via either brute force generation of mutational variants or library selections, which are only able to scan through a fraction of the available mutational landscape.

## Materials and Methods

### Model structure of EnvZ/Spo0F complex

The detailed crystal structure of a TCS complex was first obtained for HK853/RR468 of *Thermatoga maritima* (Casino et al., 2009), elucidating the binding interface between HK and RR partners. It was subsequently shown (Schug et al., 2009) that TCS complexes could be computationally predicted using highly coevolving interprotein (HK/RR) residue pairs as docking constraints for molecular dynamics simulations. In this present work, the computationally predicted structure for the KinA/Spo0F complex (*B. subtilis*) (Cheng et al., 2014) is used as a model for selecting mutations to stabilize the EnvZ/Spo0F complex. This predicted complex is composed of a representative structure for the Spo0F monomer, obtained from crystallography (PDB ID: 1PEY) (Mukhopadhyay et al., 2004), and a representative structure for the KinA homodimer, obtained from homology modeling using I-TASSER (Zhang, 2008). The sequence of EnvZ is threaded into the template structure of KinA.

### Direct Coupling Analysis (DCA)

Multiple-sequence alignments (MSA) of the HK (PF00512) and RR (PF00072) protein families are collected from Pfam (Finn, Bateman et al., 2014) (Version 28). The HK and RR aligned sequences are then concatenated based on genomic adjacency (Skerker, Prasol et al., 2005, Yamamoto, Hirao et al., 2005). The concatenated sequence of amino acids for a TCS partner pair, 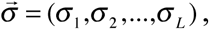 is represented as a vector of length *L*=172 where amino acids 1 to 64 and 65 to 172 belong to the HK and RR, respectively. Additional details of the TCS partners used to parameterize the coevolutionary model can be found in a previous publication (Cheng et al., 2016).

Methods such as Direct Coupling Analysis (DCA) (Ekeberg et al., 2013, Morcos et al., 2011, Weigt et al., 2009) infer a probabilistic model, 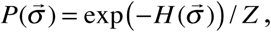 for the selection of the sequence data, 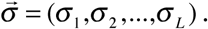 The approach adopted in this study uses pseudolikelihood maximization (Ekeberg et al., 2013) to infer the statistical couplings, *J*_*ij*_, and local fields, *h*_*i*_, of a Potts model, 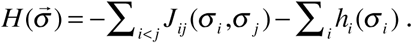

### Construction of the TCS coevolutionary landscape

Focusing on the interprotein couplings between the HK and RR residues, a proxy for signal transfer efficiency between TCS proteins in constructed:

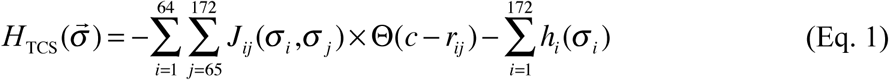

where 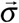 is the concatenated sequence of wild-type EnvZ (HK) and wild-type or mutant Spo0F (RR), the double summation is taken between all interprotein residue pairs (i.e., residues 1 to 64 and 65 to 172 belonging to the HK and RR, respectively), Θ is a Heaviside step function, *c* is the a cutoff distance of 16Å, and *r*_*ij*_ is the minimum distance between residues *i* and *j* in the representative structure. Mutational changes in Eq. 1 are expressed as 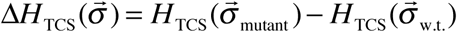 between a mutant sequence, 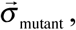 and a wild-type sequence, 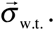 Once again, the sequence 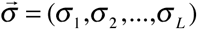 is a concatenated sequence of the HK EnvZ and the RR Spo0F, where only single-site mutations are made to the Spo0F in this present work. The coevolutionary landscape (Eq. 1) was previously published (Cheng et al., 2016), and is available online for download (http://utdallas.edu/~faruckm/PublicationDatasets.html).

It has previously been shown that coevolutionary landscapes can also be used to identify TCS partner interactions (Bitbol, Dwyer et al., 2016, Cheng et al., 2016), i.e., which HKs and RRs preferentially interact. These approaches are consistent with earlier approaches that used information-based scores (Boyd, Cheng et al., 2016, Cheng et al., 2014, Procaccini, Lunt et al., 2011).

### Direct Information (DI)

Coevolution between residue pairs *i* and *j* can be quantified using the Direct Information (DI) (dos Santos, Morcos et al., 2015, Morcos, Jana et al., 2013, Morcos et al., 2011, Weigt et al., 2009):

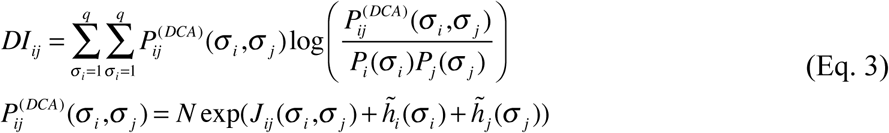

where 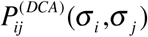 is the inferred pair distribution between residues *i* and *j* with amino acids *σ*_*i*_ and *σ*_*j*_, respectively; *N* is the normalization; and 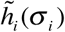 and 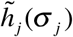 are chosen such that 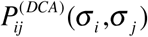 satisfies the marginalization conditions, 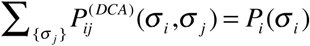 and 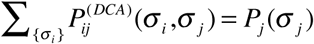 (dos Santos et al., 2015, Morcos et al., 2013, Morcos et al., 2011, Weigt et al., 2009). The DI (a Kullback-Leibler divergence) quantifies the informational entropy difference between the inferred pair distribution, 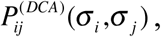 with respect to a null model lacking pairwise correlations, *P*_*i*_(*σ*_*i*_)*P*_*j*_(*σ*_*j*_).

### Protein Purification

All genes were purchased from Genescript. EnvZ was cloned into a pET-32b vector using restriction sites MscI and NcoI. Spo0F was cloned into a pET-20b(+) vector using restriction site NdeI and XhoI. Spo0F and EnvZ were transferred into BL21(DE3)pLysS and C43 competent cells, respectively, and grown in LB medium to an OD of 0.6. Protein expression was induced with the addition of 1mM IPTG for 4-5 hours at 37°C. Cells were harvested by centrifugation and resuspended in lysis buffer. Spo0F was sonicated and the supernatant was filtered through a 20 kDa filter before being loaded onto a Q-column. EnvZ was sonicated and purified with a (His)_6_-tag using a Nickel column as described (Laub, Biondi et al., 2007). In each case, fractions containing protein were pooled together, concentrated and dialyzed against phosphorylation assay buffer (see buffer conditions below). Protein purity was evaluated with SDS page.

### Phosphotransfer experiments

The phosphotransfer between EnvZ and Spo0F was measured using a radiolabeled ATP phosphotransfer assay. EnvZ and Spo0F were separately equilibrated in phosphorylation assay buffer (10 mM HEPES, 50 mM KCl, 10 mM MgCl_2_ and 0.1 mM EDTA). 100 mM ATP and 5 μCi [γ^32^P]ATP (6000 Ci/mmol) was added to the EnvZ sample to allow the autophosphorylation reaction to reach equilibrium. Equimolar amounts of phosphorylated EnvZ and Spo0F were then combined to initiate the phosphotransfer reaction. The reactions were quenched with 4x SDS Page loading buffer after 1-5 minutes, loaded on a SDS poly-acrylamide gel, run at 100 V for 1.5 hours and set to dry for 16 hours. The dried gel was exposed to film for times ranging from 10-60 minutes depending on activity, and individual protein bands corresponding to phosphorylated Spo0F were quantitated on the ^32^P channel in liquid scintillant. Reaction velocities for mutants were then calculated and expressed as a ratio compared to the wild-type enzyme.

### Thermal stability through Differential Scanning Calorimetry (DSC) measurements

To verify if the introduced point mutations have an effect on the overall protein stability, Differential Scanning Calorimetry (DSC) measurements were performed using a Microcal VP-Capilllary DSC Instrument and scanned from 20 − 100°C. DSC measures the heat change associated with thermal unfolding at a constant rate, i.e., the thermal transition midpoint (T_m_) is obtained together with the change in enthalpy (Δ*H*_DSC_) upon unfolding of the protein. Data analyses were performed using the MicroCal Origin Software, and the main transition representing the unfolding curve of monomeric Spo0F is plotted in Figure 3. Data were collected data at a 90 deg/hr scan rate on protein concentration ca. 1mg/ml.

## Acknowledgments

We would like to thank Brandon Aubol, Josh Chan and Kendra Hailey for helpful discussions.

## Author contributions

RRC, EH, NT, and FM created the collaboration. RRC, EH, NT, FM, HL, PAJ, and JNO contributed ideas to the design of the research, analyzed the data, and wrote the manuscript. RRC constructed the computational model, selected the choice of mutations, and generated the predictions. EH performed the DSC experiments and generated the mutant Spo0F proteins. NT performed the *in vitro* phosphotransfer measurements.

## Data and materials availability

The coevolutionary landscape (Eq. 1) was previously published (Cheng et al., 2016), and is available online for download (http://utdallas.edu/~faruckm/PublicationDatasets.html).

